# Bacterial Schlafens mediate anti-phage defense

**DOI:** 10.1101/2025.07.24.666596

**Authors:** Veronica Perez Taboada, Yimo Wu, Riley Cassidy, Kirill E. Medvedev, Luuk Loeff, Anna Nemudraia, Artem Nemudryi

## Abstract

Human Schlafen proteins restrict viral replication by cleaving tRNA, thereby suppressing protein synthesis. Although the ribonuclease domain of Schlafen proteins is conserved across all domains of life, its function in prokaryotes has remained unclear. Here, we show that prokaryotic Schlafen nucleases (pSlfns) are widespread antiviral effectors that protect bacteria from phages. These nucleases are fused to diverse protein domains that sense phage infection. We focus on a system where Schlafen nuclease is fused to a previously unknown immunoglobulin-like sensor domain and demonstrate that it recognizes T5-like phage tail assembly chaperones and cleaves both bacterial and viral tRNA, triggering abortive infection. Our findings redefine Schlafens as an ancient, mechanistically conserved family of immune effectors, revealing the deep evolutionary origin of tRNA-targeting antiviral immunity in humans.

---

The mammalian Schlafen (Slfn) protein family was first identified in 1998 and named for its ability to induce growth arrest in murine thymocytes (“Schlafen” is German for “sleep”)^1^. Later work found that mammalian Slfn proteins share a conserved N-terminal endoribonuclease domain that suppresses protein synthesis by cleaving tRNA and rRNA^2-6^. Several Slfn members act as interferon-inducible antiviral factors^7,8^. In particular, human SLFN11 restricts diverse viruses, including human immunodeficiency virus 1 (HIV-1)^5^, human cytomegalovirus (HCMV)^9^, and positive-strand RNA flaviviruses, like Zika and Dengue viruses^10^. Human sterile alpha motif domain-containing protein 9 (SAMD9) and its paralog, SAMD9-like protein (SAMD9L), similarly use a Slfn-like tRNase domain to inhibit translation during poxviral and lentiviral infections^11,12^.

The Slfn ribonuclease domain is highly conserved in animals and, in addition to antiviral defense, has been linked to suppressing transposable elements in roundworms^13^. This recurring recruitment of the Slfn ribonuclease domain for immune functions suggests deep evolutionary roots of its role in restricting genetic parasites^8,13,14^. Slfn domains are found in prokaryotic genomes, and a few instances have been reported to associate with anti-phage genes, hinting at a potential immune function^15-17^. However, direct experimental evidence for the antiviral role of prokaryotic Schlafens (pSlfns) has been lacking, and their molecular mechanisms have not been determined.

In this work, we use a combination of computational, functional, and biochemical assays to demonstrate that pSlfn domains are anti-phage effectors fused to a variety of potential phage sensors and determine the molecular mechanism of phage defense mediated by pSlfn nuclease fused to an immunoglobulin-like domain. We show that pSlfn-mediated tRNA cleavage triggers growth arrest in response to phage infection, revealing evolutionary, functional, and mechanistic conservation with human Slfn-mediated antiviral immunity.

## Bacterial Schlafen proteins mediate anti-phage defense

Human SLFN11 protein has an N-terminal ribonuclease (RNase) domain, a linker domain, and a C-terminal DNA/RNA helicase domain^4^. The N-terminal domain consists of two lobes (N- and C-lobe), where the C-lobe contains the ribonuclease active site with a catalytic Ex_4_ExK motif that is essential for tRNA cleavage (Fig. 1a)^4^. To identify prokaryotic Slfn homologs, we searched for Slfn nuclease domains in reference prokaryotic genomes and identified 5,930 unique (9,937 total) protein sequences across 4 archaeal and 34 bacterial phyla (Fig. 1a, Supplementary Fig. 1a, and Supplementary Table 1). Subsequent multiple sequence alignments show a highly conserved Ex_4_(E/D)xK sequence motif in the predicted active site, which matches the catalytic residues of SLFN11, suggesting a conserved enzymatic mechanism (Fig. 1a; Supplementary Fig. 1b).

**Fig 1.**
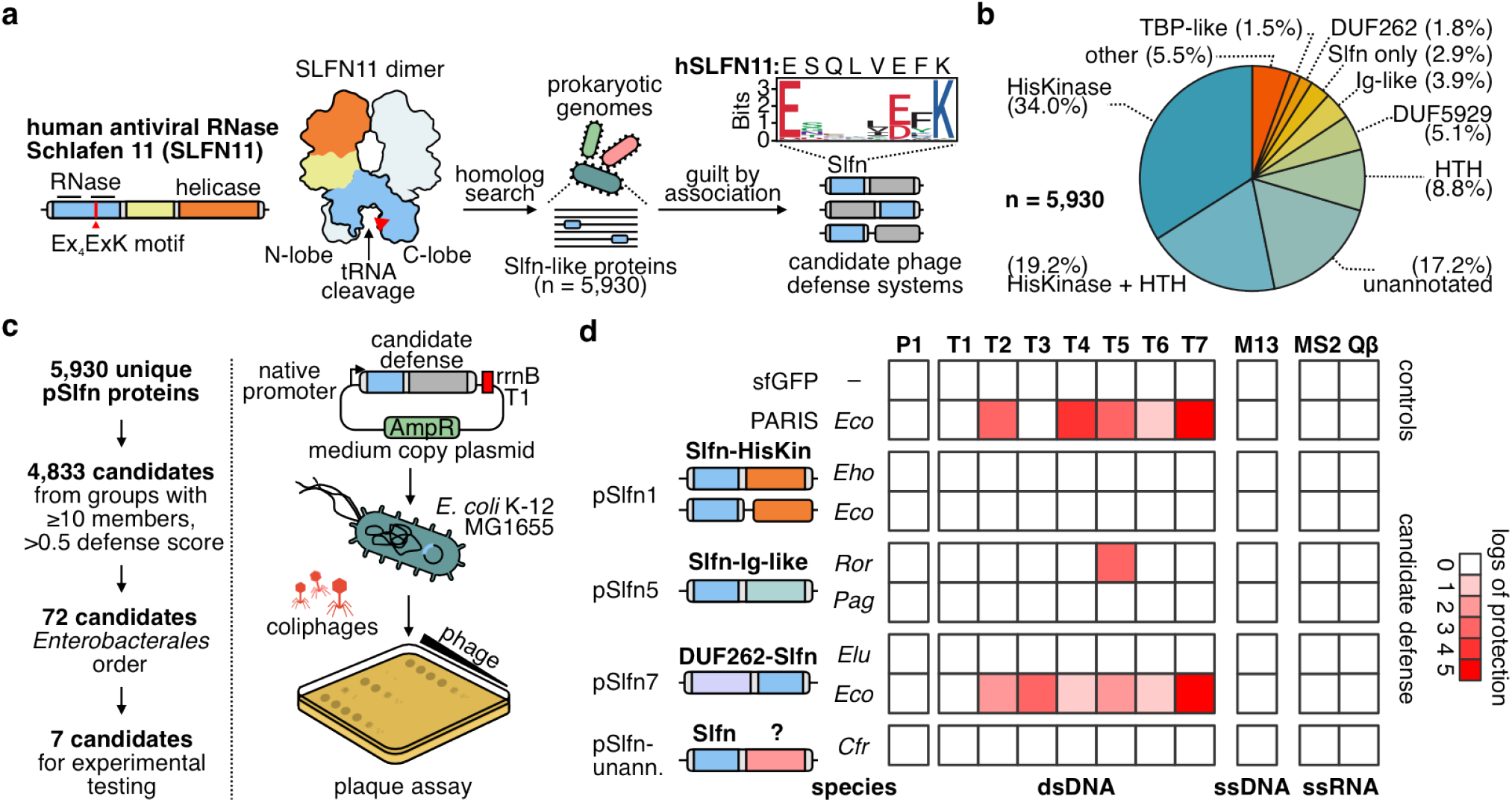
Schlafen proteins mediate phage defense in bacteria. **a**, Computational approach to identify prokaryotic homologs of Schlafen nucleases (Slfn) with predicted roles in phage defense. This search found 5,930 unique Slfn protein sequences (100% CD-HIT cut-off). Weblogo plot shows conserved Ex_4_(E/D)xK motif in prokaryotic Slfn domains. **b**, Annotation of domains fused to Slfn in the 5,930 identified proteins. **c**, Experimental approach to test for immune activities of candidate pSlfn phage defense. **d**, Immune activities of tested Slfn proteins (top-to-bottom) against a panel of phages that infect *E. coli* (left-to-right). *Eco* – *Escherichia coli, Eho* - *Enterobacter hormaechei, Ror* - *Raoultella ornithinolytica, Pag* - *Pantoea agglomerans, Elu* - *Enterobacter ludwigii, Cfr* - *Citrobacter freundii*.

Prokaryotic defense genes frequently cluster in the host genome, forming genetic neighborhoods known as defense islands^15,18^. To determine whether pSlfn proteins reside in defense islands, we performed a genetic neighborhood analysis and found that 69.6% of identified pSlfn genes are located within a +/− 10 kb distance to at least one known defense gene (Supplementary Fig. 1a, and Supplementary Table 1). On average, identified pSlfn genes co-localize with 1.9 defense genes, which is comparable to AriB (2.74), the nuclease effector of the phage anti-restriction-induced system (PARIS)^19,20^, and approximately tenfold higher than the housekeeping genes rpL3 (0.13) and L1p (0.19) (Supplementary Fig. 1c). This genetic association suggests that pSlfns play a role in anti-phage defense. Notably, 30.4% of identified pSlfn genes were not associated with defense genes, suggesting that in some instances, pSlfn domains might have non-immune functions or that these pSlfn genes are in defense hotspots that have not been annotated yet^20,21^.

Phage defense systems often function through sensors that detect infection and effectors that execute the immune response. These functional modules can be encoded within a single gene or split across multiple genes^22,23^. We found that 97.1% of identified pSlfn domains are fused to other protein domains, where pSlfns most likely act as effectors, while identified fused domains may function as sensors (Fig. 1b). To classify the diversity of domain architectures among pSlfn proteins, we used a combination of sequence and structural homology-based computational methods, identifying 55 distinct domain compositions, which we designated pSlfn1 through pSlfn55 (Supplementary Table 2). The most frequent domain compositions include fusions of pSlfn to histidine kinase (HisKinase, 34.0%), HisKinase and helix-turn-helix domains (HisKinase-HTH, 19.2%), HTH domains (8.8%), DUF5929 (5.1%), domains with immunoglobulin-like fold (Ig-like, 3.9%), DUF262 (1.8%), and TBP-like domains (1.5%) (Fig. 1b). Of 5,930 identified pSlfns, 1,020 (17.2%) are fused to protein domains that show no significant homology to the domains annotated in the Pfam and ECOD (Evolutionary Classification of protein Domains) databases^24,25^. We further grouped these proteins into 558 sequence similarity clusters (Supplementary Table 2). This diversity of domain compositions in pSlfn proteins likely represents combinations of the core Slfn effector module with various sensors that respond to different cues and regulate its nuclease activity.

To test whether pSlfn genes confer anti-phage defense, we expressed seven representatives from *Enterobacterales* in *Escherichia coli* K-12 strain MG1655 under their native promoters (Supplementary Fig. 1d; Supplementary Table 3), and challenged the cells with a panel of diverse *E. coli* phages (Fig. 1c,d). As a positive control, we used a plasmid encoding the *E. coli* B185 PARIS immune system^20^, while a plasmid expressing sfGFP served as a negative control. In the tested subset, two pSlfn genes provided robust defense against T-phages (Fig. 1d; Supplementary Fig. 1e). We found that DUF262-Slfn fusion from *E. coli* UC4224 protects bacteria from a broad range of phages, with the most potent activity being against T3 and T7 phages. In contrast, Slfn protein from *Raoultella ornithinolytica* (*Ror*Slfn5) protected *E. coli* only against T5 phage. Overall, these data demonstrate that prokaryotic Schlafen proteins with distinct domain architectures protect bacteria from phage infection.

## T5 tail assembly chaperone triggers *Ror*Slfn5-mediated abortive infection

To elucidate the mechanisms that underpin the immune function of pSlfn proteins, we focused on *Ror*Slfn5 defense. We challenged MG1655 cells with T5 phage at different multiplicities of infection (MOIs) and measured cell growth. At low MOIs (0.01 and 0.1), cells expressing *Ror*Slfn5 survived the infection compared to cells without the defense. In contrast, infections at high MOIs (5 and 10) resulted in bacterial cultures collapsing approximately sixty minutes post-infection, which is similar to the lysis time of T5 phage^26^ (Fig. 2a, b). Mutations in the predicted Slfn catalytic motif Ex_4_DxK (E15A & D20A) abolished defense, indicating that the nuclease activity of pSlfn5 is required for its anti-phage function (Fig. 2c). At an MOI of 10, at which nearly all (>99.9%) of cells are infected at once, cells expressing *Ror*Slfn5 collapsed approximately 10 minutes faster than cells without the defense (Fig. 2d). Furthermore, T5 phage infection (MOI = 10) of cells with *Ror*Slfn5 produced 2.5-fold less plaque-forming units (PFU) after one viral replication cycle (60 min) compared to cells without the defense (*p* = 0.008; Fig. 2e). The premature culture collapse (Fig. 2d) and reduced phage propagation (Fig. 2e) indicate that activation of the pSlfn5 defense triggers nan abortive infection phenotype^27^.

**Fig 2.**
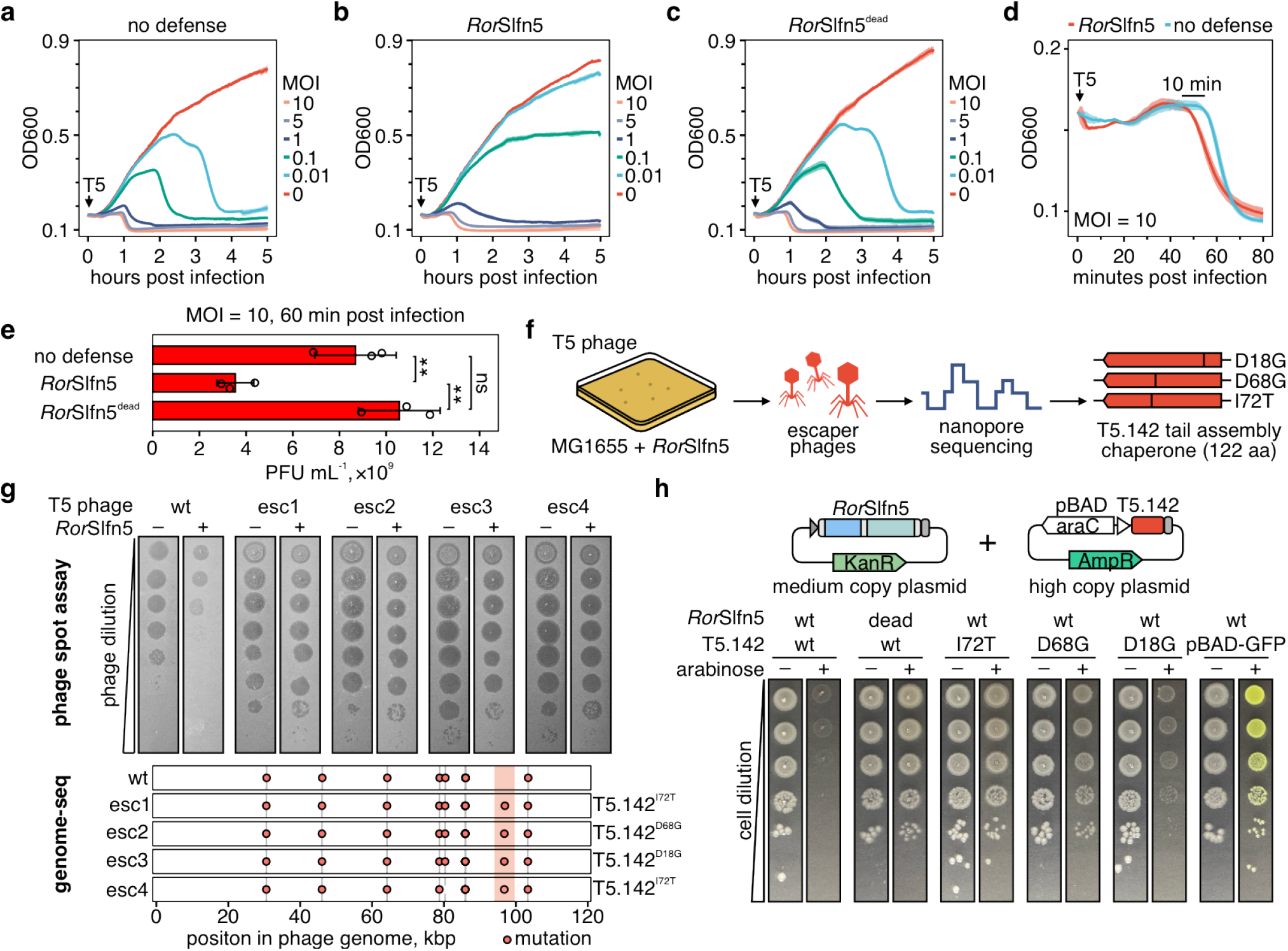
Phage tail assembly chaperone triggers pSlfn5-mediated abortive infection phenotype. **a**, Growth kinetics of *E. coli* K-12 MG1655 cells without the defense, **b**, with *Ror*Slfn5, or **c**, nuclease-dead *Ror*Slfn5^E15A, D20A^ mutant following T5 bacteriophage infection at various multiplicities of infection (MOIs). Data is shown as the mean of four replicates (center line) *±* S.D. (ribbon). **d**, Comparison of growth kinetics during the first 80 minutes post-infection. **e**, Quantification of plaque-forming units (PFU) in cellular supernatants after 60 minutes of T5 phage infection. Data is shown as the mean of three biological replicates *±* S.D. One-way ANOVA with post hoc Tukey HSD test was used to compare the experimental groups. ** – *p <* 0.01; ns – non-significant. **f**, An experimental approach to identify viral triggers of *Ror*Slfn5. **g**, *Top*: plaque assays with *Ror*Slfn5-escaping T5 phages (esc1-esc4). *Bottom*: mapping of mutations in T5 phage escapers compared to a reference phage genome. Vertical lines indicate mutations found in the lab T5 phage stock compared to the reference NCBI sequence. **h**, Toxicity assay in MG1655 cells co-transformed with *Ror*Slfn5 plasmid and a plasmid for arabinose-inducible expression of T5.142 protein. Ten-fold serial dilutions were spotted on agar plates with 0.2% glucose (-) or 0.2% arabinose (+). pBAD-GFP was used as a control for arabinose-dependent induction.

To determine the viral trigger for the *Ror*Slfn5 defense, we isolated T5 phages that escaped the immunity and sequenced their genomes (Fig. 2f). Four viral clones that we sequenced contained three distinct missense mutations (i.e., D18G, D68G, and I72T) in the T5.142 gene, which encodes for the phage tail assembly chaperone (NCBI accession: YP_006970.1) (Fig. 2g, Supplementary Table 5)^28^.

To test if the T5.142 protein triggers *Ror*Slfn5 defense, we cloned the viral gene and its escape variants into high-copy plasmid vectors under the control of the arabinose-inducible promoter (pBAD; Fig. 2h). Expression of T5.142 in cells co-transformed with *Ror*Slfn5 plasmid resulted in toxicity, demonstrating that this viral protein is sufficient to trigger the defense. In contrast, cells expressing a catalytic mutant of *Ror*Slfn5 (*Ror*Slfn5^E15A, D20A^) demonstrated no growth defect compared to the uninduced control cells, indicating that the toxicity is mediated by T5.142-triggered nuclease activity of *Ror*Slfn5. In addition, escape mutations in the viral T5.142 protein rescued toxicity. While the T5.142^D18G^ variant reduces toxicity, it does not completely abolish it, suggesting that in our experimental setup, it is less effective at evading *Ror*Slfn5 defense than the D68G and I72T mutations.

## The immunoglobulin-like domain of pSlfn5 is a phage sensor

Homologs of pSlfn5 from *Pantoea agglomerans* (*Pag*Slfn5; 39.6% sequence identity to *Ror*Slfn5) and *Serratia fonticola* (*Sfo*Slfn5; 79.6% sequence identity to *Ror*Slfn5) did not protect *E. coli* from T5 phage or any other phage tested (Supplementary Fig. 2a). The sequence divergence between these proteins is primarily driven by their C-terminal domains, while the N-terminal Slfn nuclease domains are more conserved (Supplementary Fig. 3a). AlphaFold-predicted structure of *Ror*Slfn5 suggests that its C-terminal domain has an immunoglobulin-like (Ig-like) *β*-sandwich fold with structural similarity to human integrin ectodomains (Supplementary Fg. 3b-c; Supplementary Table 4). These observations led us to hypothesize that the Ig-like domains of pSlfn5 proteins function as sensor modules that recognize viral proteins, and thereby *Pag*Slfn5 and *Sfo*Slfn5 might recognize T5.142 homologs from other phages.

To test this, we synthesized genes encoding six representative homologs of the T5 phage tail assembly chaperone (Fig. 3a). In addition to T5.142, *Ror*Slfn5 was triggered by expression of homologs from *Salmonella* phage S114, *Pantoea* phage vB_PagS_AAS21, and an unclassified phage from a human gut metagenome (Fig. 3b, Supplementary Fig. 2b-d). Consistent with the lack of defense against the T5 phage, *Pag*Slfn5 and *Sfo*Slfn5 were not activated by expression of T5.142, while the tail assembly chaperone from the human gut metagenome triggered all tested pSlfn5 homologs (Fig. 3b). None of the identified triggers caused toxicity when expressed alone or co-expressed with a nuclease-dead *Ror*Slfn5^E15A, D20A^ mutant, confirming that toxicity is mediated by the nuclease activity of pSlfn5 (Supplementary Fig. 2e, f). Together, these findings demonstrate that pSlfn5 proteins recognize tail assembly chaperones from phages in the *Demerecviridae* family, with different pSlfn5 homologs exhibiting distinct trigger specificities.

**Fig 3.**
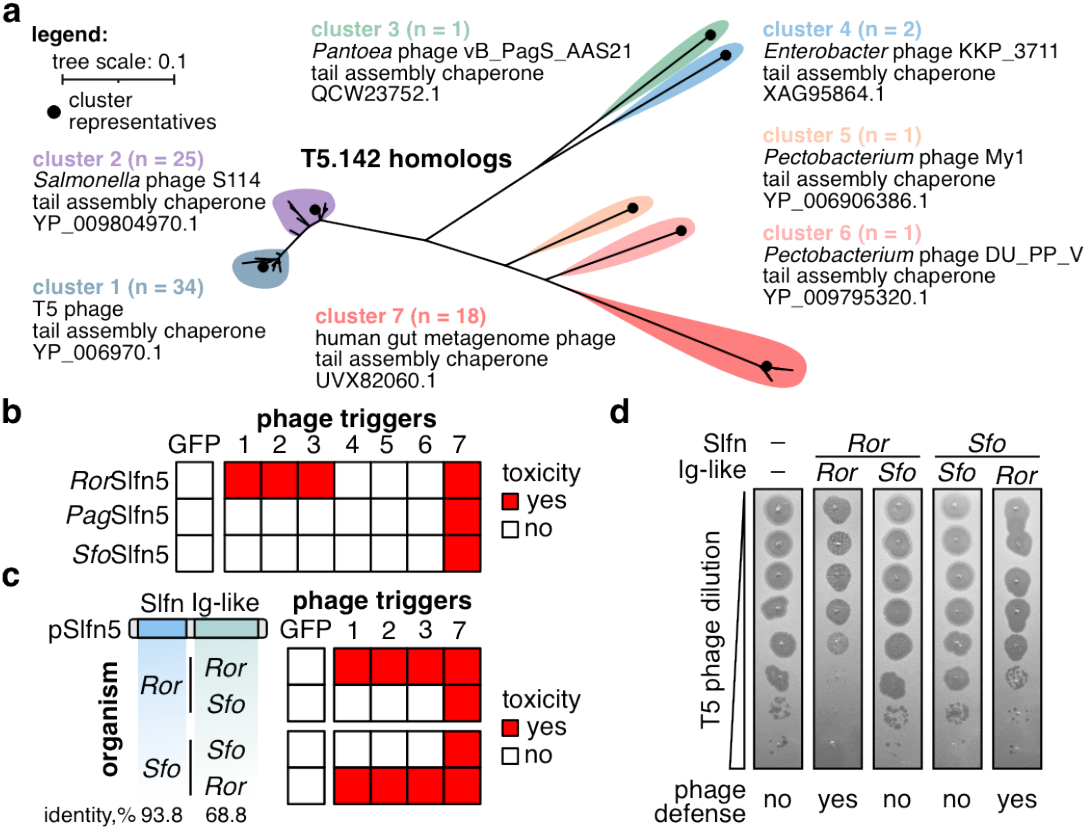
The Ig-like domain mediates the specificity of pSlfn5 immunity. **a**, Maximum-likelihood phylogenetic tree of T5.142 homologs. Selected representatives are listed for each homolog cluster and indicated with black dots. **b**, Summary of toxicity assays in MG1655 cells co-transformed with pSlfn5 (top-to-bottom) and T5.142 homologs from various phages (left-to-right). Numbers correspond to homolog clusters in (a). **c-d**, Toxicity (c) and plaque assay (d) with chimeric pSlfn5 proteins with swapped Ig-like domains.

To confirm the sensor function of Ig-like domains in pSlfn5 defense, we created chimeric proteins by swapping the Ig-like domains between *Ror*Slfn5 and *Sfo*Slfn5. Cellular toxicity assays showed that phage trigger specificity was transferred along with the sensor domain (Fig. 3c, Supplementary Fig. 4). Furthermore, the chimeric protein containing the nuclease domain of *Sfo*Slfn5 and the *Ror*Slfn5 sensor gained protection against T5 phage, whereas changing the Ig-like domain of *Ror*Slfn5 for that of *Sfo*Slfn5 abolished defense against T5 (Fig. 3d). Overall, these data demonstrate that Ig-like domain dictates phage specificity and functions as a sensor module of the pSlfn5 defense system.

## *Ror*Slfn5 is a phage-activated tRNase

After identifying the viral cue that triggers pSlfn5 defense, we next investigated the mechanism of defense. Co-expression of *Ror*Slfn5 with its viral trigger led to growth arrest ~30 minutes after inducing T5.142 expression (Fig. 4a). This timing mirrors the kinetics of GFP expression under the same promoter, which became detectable 30 minutes post-induction (Fig. 4b). Control cells co-expressing *Ror*Slfn5 and GFP or *Ror*Slfn5^E15A, D20A^ and T5.142 showed no growth arrest, demonstrating that it is mediated by T5.142-triggered nuclease activity of the pSlfn domain (Supplementary Fig. 5a, b).

**Fig 4.**
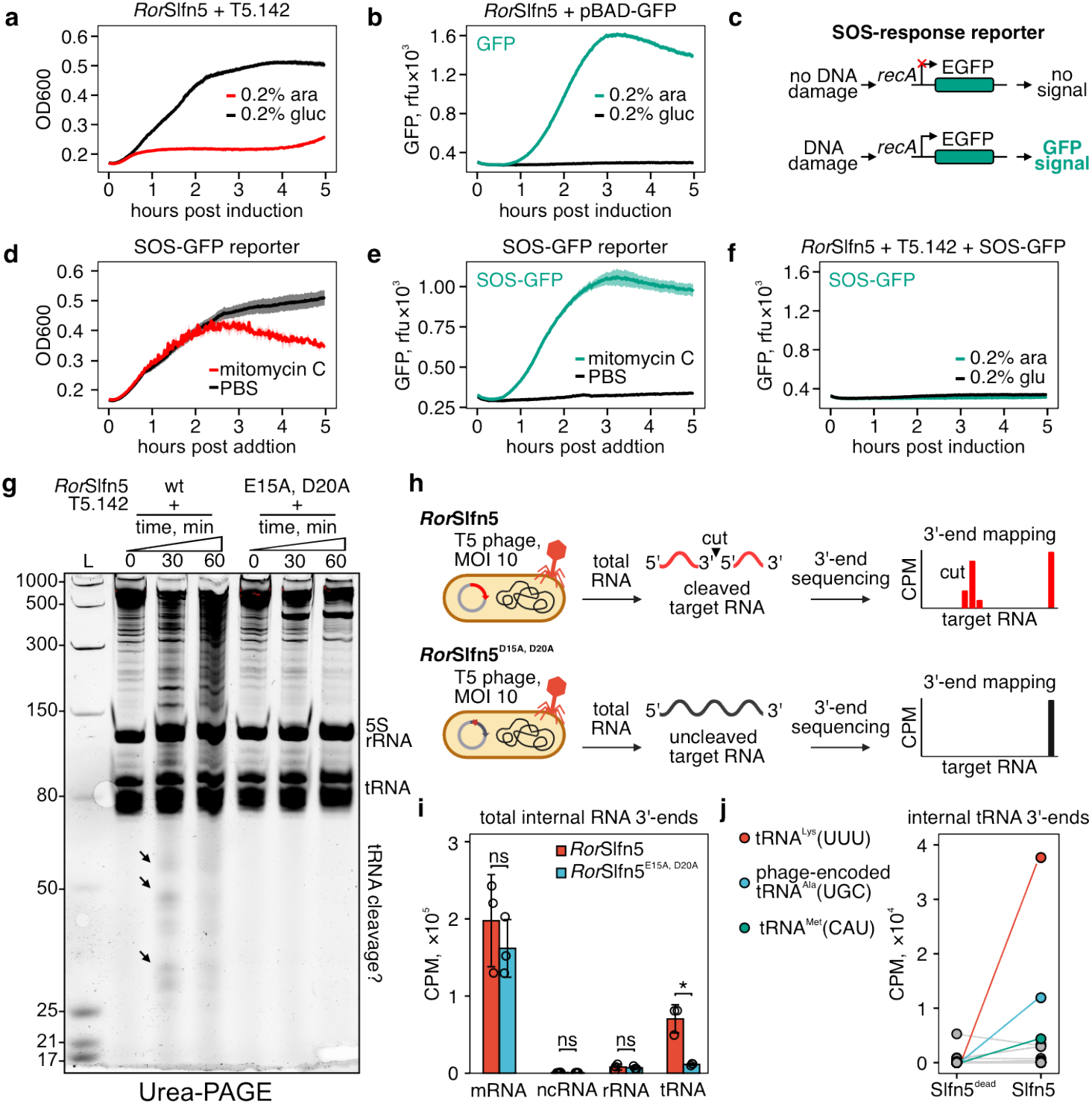
*Ror*Slfn5 cleaves tRNA upon phage infection. **a**, Growth kinetics of MG1655 cells constitutively expressing *Ror*Slfn5 with (0.2% arabinose [ara]) and without (0.2% glucose [gluc]) induction of T5.142 expression. **b**, Kinetics of GFP expression after induction of the pBAD promoter with arabinose. **c**, Schematics of reporter assay for detecting SOS-response upon DNA damage. **d-e**, Growth kinetics (**d**) and SOS-response reporter assay (**e**) of MG1655 treated with 100 nM mitomycin C or PBS. **f**, SOS-response reporter assay with *Ror*Slfn5-expressing cells after induction of T5.142 expression. Kinetic assays in **a-f** were performed in four biological replicates. Data is shown as mean (center line) *±* S.D. (ribbon). **g**, Urea-PAGE of total RNA extracted from *RorSlfn* or *RorSlfn*^E15A, D20A^-expressing MG1655 at various timepoints after induction of T5.142 expression. **h**, Schematics of the 3’-end RNA sequencing approach used to map *Ror*Slfn5-mediated RNA cleavage in T5 phage-infected cells. **i**, Total abundance of non-native (internal) 3’-ends in different RNA types in T5 phage-infected cells expressing *Ror*Slfn5 or its nuclease-dead version (E15A and D20A mutation). Data is shown as the mean of three biological replicates *±*S.D. Means were compared using a two-sided Welch’s t-test. **\*** *p <* 0.05, ns – non-significant. **j**, Total abundance of internal 3’-ends in bacterial and viral tRNAs. Data is shown as the mean of three replicates.

To determine if *Ror*Slfn5 targets DNA, we tested whether T5.142-triggered *Ror*Slfn5 nuclease activity causes bacterial DNA damage. We used a fluorescent reporter plasmid (pSOS-GFP) that encoded a *gfp* gene under control of *recA* promoter, which is upregulated in the bacterial SOS response to DNA damage (Fig. 4c). Treatment of MG1655 with mitomycin C, a DNA-damaging agent, induced a strong fluorescent signal, which was absent in PBS-treated cells (Fig. 4d, e). In contrast, cells co-transformed with *Ror*Slfn5, T5.142, and pSOS-GFP did not show increased fluorescence, suggesting that *Ror*Slfn5 activation does not cause DNA damage and the SOS response (Fig. 4f; Supplementary Fig. 5c, d). These results indicate that *Ror*Slfn5 does not act on DNA and likely retains RNase activity as its primary mechanism of toxicity, consistent with the function of human SLFN proteins^3,4^.

To test if *Ror*Slfn5 targets RNA, we co-transformed MG1655 cells with plasmids encoding the T5.142 trigger and either wild-type *Ror*Slfn5 or a catalytically inactive mutant *Ror*Slfn5^E15A, D20A^. We extracted total RNA from cells at 0, 30, and 60 minutes post-T5.142 induction. Urea-PAGE analysis of extracted RNA identified RNA fragments between 25 and 50 nt in size that appeared at 30- and 60-minute post-induction in cells expressing *Ror*Slfn5 but not the catalytically inactivated mutant, demonstrating that T5.142 expression triggers *Ror*Slfn5-mediated RNA cleavage (Fig. 4g).

To map RNA cleavage induced by *Ror*Slfn5, we performed RNA 3’-end sequencing (Fig. 4h). To validate this approach, we analyzed RNA from *E. coli* expressing the PARIS immune system. RNA sequencing (RNA-seq) revealed a significant enrichment of internal 3’-ends in tRNA^Lys(UUU)^ (FDR = 0.002) in T5-infected cells expressing the active PARIS defense compared to cells expressing the inactivated PARIS system (AriB^E26A^ mutant; Supplementary Fig. 6a-c). This result agrees with previous work^19,29^ demonstrating that activation of PARIS immunity triggers AriB-mediated cleavage of tRNA^Lys(UUU)^.

Next, we applied the same RNA-seq strategy to T5 phage-infected MG1655 cells expressing *Ror*Slfn5 or its nuclease-dead mutant (Fig. 4h, Supplementary Fig. 6g). This analysis found no significant difference in the abundance of internal 3’-ends in messenger RNA (mRNA), non-coding RNA (ncRNA), or ribosomal RNA (rRNA). In contrast, we observed a 6.2-fold increase in internal 3’-ends in tRNA of cells expressing wild-type *Ror*Slfn5 (*p* = 0.03; Fig. 4i). Analysis of tRNA-mapping reads identified significant enrichment of total internal 3’-end counts in tRNA^Lys(UUU)^ (FDR = 0.01), phage-encoded tRNA^Ala(UGC)^ (FDR = 0.04), and tRNA^Met(CAU)^ (FDR = 0.02) in cells expressing wild-type *Ror*Slfn5 compared to the nuclease-dead mutant (Slfn5^dead^; Fig. 4j). Position-specific analysis confirmed these enrichments and identified additional tRNAs with significant cleavage signatures, including tRNA^Val(UAC)^, tRNA^Phe(GAA)^, and tRNA^Ile(CAU)^ (FDR < 0.05; Supplementary Fig. 6h). Together, these results indicated that *Ror*Slfn5 preferentially cleaves tRNA upon phage infection, with tRNA^Lys(UUU)^ fragments being most abundant in the cell.

## Phage-activated *Ror*Slfn5 cleaves the tRNA anticodon arm

We next sought to map *Ror*Slfn5-mediated tRNA cleavage sites at the single-nucleotide resolution. Mapping of 3’-ends in tRNA^Lys(UUU)^ identified three highly enriched positions within the anticodon arm (29, 30, and 32), while the 5’-end mapping showed peaks at positions 39 and 41 (Fig. 5a, b). Correspondingly, sequence coverage across the entire anticodon loop of tRNA^Lys(UUU)^ was absent, indicating complete removal of this region (Fig. 5c). A similar sequence coverage gap was observed between the 3’- and 5’-ends of tRNA^Met(CAU)^ fragments (Supplementary Fig. 6i-k, m). This cleavage pattern suggests that *Ror*Slfn5 may either excise the anticodon loop through two precise cuts or perform a single cut followed by trimming of the resulting fragments by host ribonucleases. Supporting this model, RNA-seq of cells expressing the PARIS immune system revealed a similar 11-nucleotide gap in the anticodon loop of tRNA^Lys(UUU)^ spanning nucleotides 29 and 41, the latter position matching the AriB cut site identified *in vitro*^19,29^. These findings support a mechanism in which cleavage is followed by trimming of the anticodon loop (Supplementary Fig. 6d-f, l).

**Fig 5.**
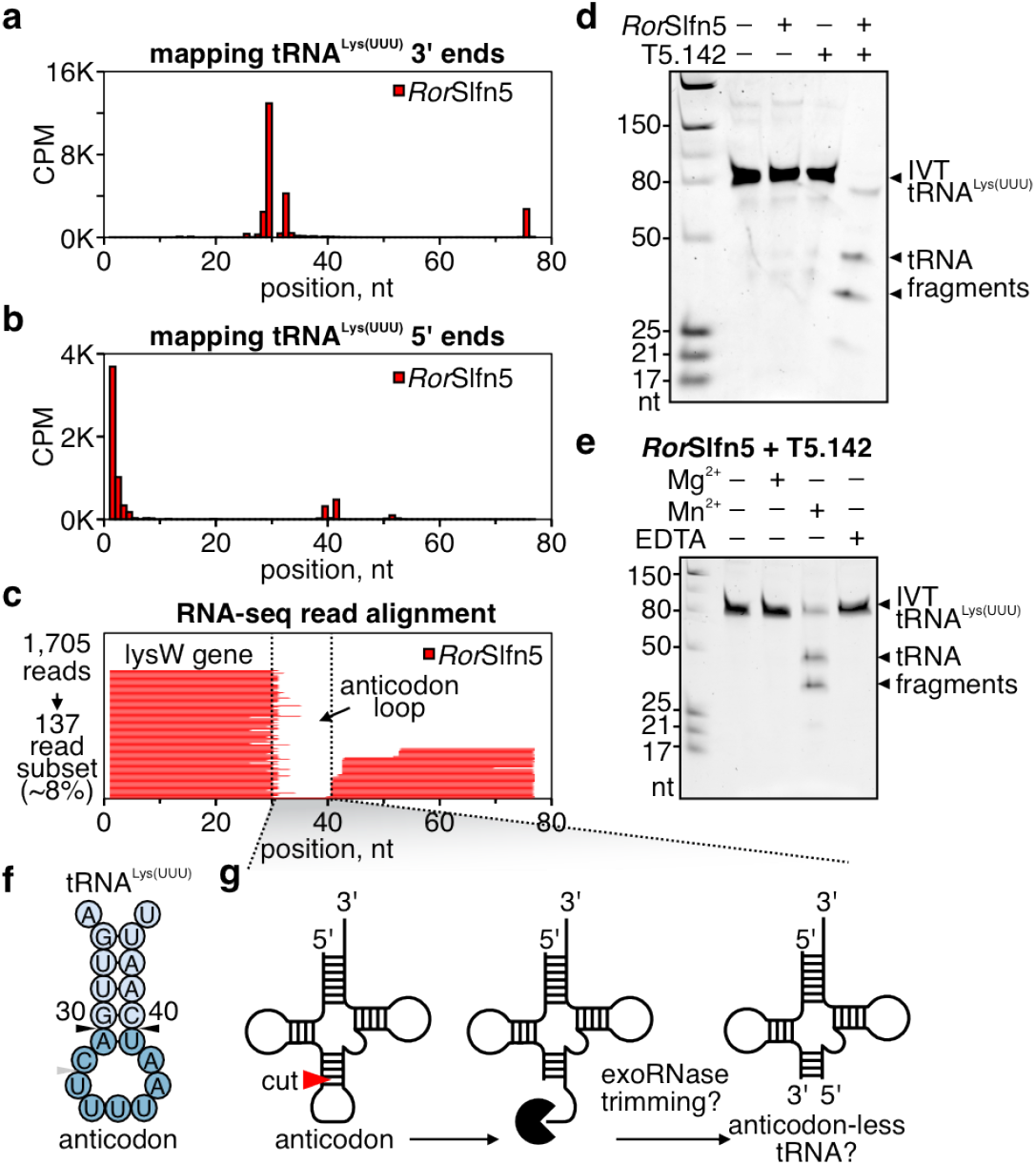
Phage-activated *Ror*Slfn5 cleaves the tRNA anticodon arm. **a, b**, Position-specific mapping of internal 3’-ends (**a**) and 5’-ends (**b**) in tRNA^Lys(UUU)^ in T5-phage-infected cells expressing *Ror*Slfn5. Data is shown as the mean of three biological replicates. **c**, Alignment of sequencing reads to the *lysW* gene of MG1655 cells. One representative replicate from three biological replicates is shown. **d**, tRNA cleavage assays with 100 nM tRNA^Lys(UUU)^ and 100 nM *Ror*Slfn5 in the presence of trigger T5.142 (2 *µ*M). **e**, Same as in (**d**) but with Mg^2+^, Mn^2+^, or no metal and EDTA. **f**, Potential *Ror*Slfn5 cut sites (black triangles) in the anticodon arm of tRNA^Lys(UUU)^, according to RNA-seq data in (**a**) and (**b**). **g**, Proposed model for *Ror*Slfn5-mediated tRNA cleavage *in vivo* and subsequent host exoribonuclease (exoRNase) trimming of the anticodon loop.

To reconstitute tRNA cleavage *in vitro*, we expressed and purified *Ror*Slfn5 and T5.142 proteins. The size-exclusion chromatography (SEC) profile of the *Ror*Slfn5 (43.1 kDa) indicates that it assembles in higher-order oligomers, while T5.142 (13.8 kDa) eluted at a size consistent with a dimer or trimer, based on gel filtration standards (Supplementary Fig. 7a,b). As Ig-like domains often mediate protein-protein interactions^30^, we hypothesized that the Ig-like domain of *Ror*Slfn5 may detect phage by directly binding T5.142. To test this hypothesis, we expressed the two proteins separately, mixed the cell lysates, and performed an affinity pull-down of tagged *Ror*Slfn5. *Ror*Slfn5 and T5.142 co-purified and co-eluted in a single SEC peak, confirming a direct interaction and suggesting that binding of T5.142 triggers the tRNase activity of *Ror*Slfn5 (Supplementary Fig. 7c).

Incubation of synthetic tRNA^Lys(UUU)^ with purified *Ror*Slfn5 and T5.142 proteins, but not either of the proteins alone, produced two major cleavage fragments sized between 25 and 50 nucleotides in length (Fig. 5d). Similar to human SLFN nucleases^3,4^, *Ror*Slfn5-mediated tRNA cleavage is manganese-dependent, while addition of magnesium did not support the reaction (Fig. 5e). Mutations in the pSlfn active site (E15A, D20A) abrogated the cleavage, confirming that the Slfn nuclease domain of *Ror*Slfn5 catalyzed tRNA cleavage (Supplementary Fig. 8a). Testing additional tRNA substrates showed that bacterial tRNA^Lys(UUU)^ and phage-encoded tRNA^Ala(UGC)^ were cleaved most efficiently, consistent with these tRNAs being the top hits in our RNA-seq data (Supplementary Fig. 8b; Fig. 4j).

*In vivo* RNA-seq data identified two candidate *Ror*Slfn5 cut sites for in tRNA^Lys(UUU)^, between nucleotides 30-31 (site 1) and 39-40 (site 2) (Fig. 5f). Cleavage of tRNA^Lys(UUU)^ *in vitro* produced two major products, consistent with a single cut. Based on the observed fragment sizes, the cleavage pattern supports cleavage at site 1 (30 and 46 nt products) over site 2 (37 and 39 nt products). Therefore, we propose that *in vivo RorSlfn5* cleaves the anticodon arm between nucleotides 30-31, followed by trimming of the 3’ tRNA fragment by cellular ribonuclease (Fig. 5g).

## Discussion

The evolutionary and functional conservation of the Schlafen ribonuclease domain suggests that this effector module belongs to an ancient ancestral immune system that predates the divergence of prokaryotes and eukaryotes^31^ (Fig. 6). In prokaryotes, this module is fused to a variety of accessory domains that likely act as phage sensors, regulating the nuclease activity in response to distinct viral cues. Most of these domains have not been previously associated with phage defense, revealing a previously unrecognized repertoire of sensing mechanisms that may extend across the broader prokaryotic immunity landscape. In this study, we show that the immunoglobulin-like sensor domain of pSlfn5 recognizes phage tail assembly chaperones, while the signals that trigger other Schlafen systems remain to be determined.

**Fig. 6.**
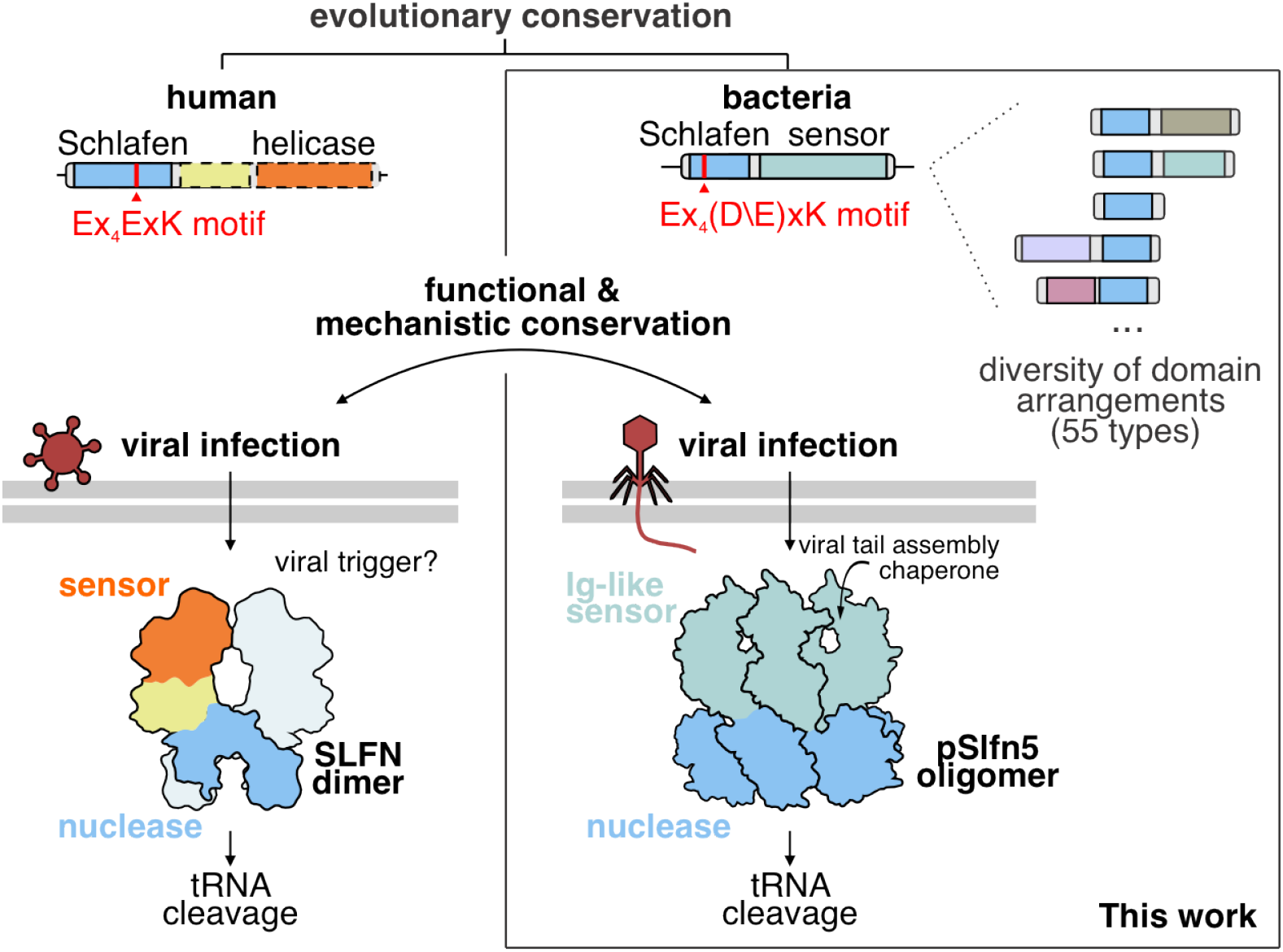
Schlafen proteins bridge innate immunity across domains of life.

The modular architecture of Schlafen proteins appears to be a deeply conserved feature, with the core ribonuclease domain fused to diverse regulatory modules in both prokaryotes and mammals. The helicase domain of human SLFN11 was recently shown to bind single-stranded DNA and activate its tRNase activity^6^, whereas the upstream triggers for SAMD9/9L remain unknown^12^. The recurring evolutionary strategy of pairing the Schlafen effector with different sensory domains underscores its remarkable success as a versatile antiviral effector.

tRNA cleavage as an antiviral strategy was first demonstrated with the discovery of the bacterial PrrC anticodon nuclease^32, 33^. Since then, several other examples of anti-phage tRNases have been described, such as AriB from the PARIS system^19^, PtuAB from Retron Ec78^34^, and ribonucleases in prophage competition elements^35^. More recently, virus-induced tRNA cleavage has also been recognized as a component of antiviral defense in mammals^3, 4, 12^. Although mechanistically similar, these shared antiviral strategies were thought to have emerged independently through convergent evolution^12^. Our findings establish a direct evolutionary link by demonstrating that homologs of human Schlafen tRNA ribonucleases function in bacterial phage defense through a tRNA cleavage mechanism that is conserved across the tree of life.

## Methods

### Plasmids, bacterial strains, and bacteriophages

The oligonucleotide primers and plasmid DNA sequences are listed in Supplementary Table 3. Candidate phage defense genes with native promoters (100-150 upstream sequence) were synthesized and cloned into a medium-copy plasmid vector (pTwist_Amp_MC) by Twist Bioscience. The plasmid expressing the *PARIS-2* immune system of *E. coli* B185 was a gift from Dr. Blake Wiedenheft^19^. Site-directed mutagenesis was used to make nucleotide substitutions in plasmid pAN248 to create the nuclease-dead variant of *Ror*Slfn5 (E15A, D20A mutations) and in plasmid pRF85 to inactivate AriB nuclease (E26A mutation) in the *PARIS-2* system. The genes encoding the T5 tail assembly chaperone (T5.142; NCBI accession: YP_006970.1) and its variants (D18G, D68G, and I72T) were PCR-amplified from genomic DNA of the T5 phage and its escape variants. Amplified genes were cloned into the pBAD vector. Homologs of T5.142 gene were synthesized by Twist Bioscience and cloned into pBAD. To perform the cell toxicity assays, the plasmid backbones of pAN248 (*Ror*Slfn5), pAN254 (*Pag*Slfn5), and pAN294 (*Sfo*Slfn5) were modified to replace the ampicillin resistance (AmpR) gene with the kanamycin resistance (KanR) gene. Plasmids pAN309 and pAN310 expressing chimeras of *RorSlfn* and *Sfo*Ig-like and *SfoSlfn* and *Ror*Ig-like were assembled using Hi-Fi (NEB, E2621). Gene fragments for pAN301 (pSOS-GFP) were synthesized by Twist Biosciences and assembled using NEBridge Golden Gate Assembly Kit (BsmBI-v2) (NEB, E1602S). For protein expression, the genes encoding for *Ror*Slfn5 and T5.142 were cloned into the pDF0118 vector backbone (Addgene #172503) in frame with a 6xHis-TwinStrep-SUMO affinity tag. All plasmid sequences were confirmed with whole-plasmid sequencing at Plasmidsaurus (https://www.plasmidsaurus.com/). All bacterial strains and phages used in this work are listed in Supplementary Table 3.

### Computational search for prokaryotic Slfn

The initial search was done using *jackhmmer* to find homologs of the human *SLFN11* nuclease domain (amino acids 1-354; NCBI: NP_001098057.1) in a database of reference proteomes^35^. The hidden Markov model (HMM) generated by *jackhmmer* was used to perform an *HMMsearch* (from HMMER 3.1b2) on a database of bacterial and archaeal amino acid sequences, created by translating RefSeq complete genomes, downloaded in July 2023, using *prodigal*^36^. The non-redundant hits from the first *HMMsearch* (e-value < 0.001; CD-HIT with `-c 0.9`) were aligned using MAFFT v7.526. A new HMM was created from the alignment using HMMbuild, and *HMMsearch* was repeated on the same database. The second round found 9,937 potential homologs (e-value < 0.001) with full-length Slfn domains (>100 aa; PF04326). After redundancy (CD-HIT with `-c 1`) was removed, a total of 5,930 protein sequences were left. Prokaryotic Slfn domains were mapped in the *HMMsearch* hits using *HMMscan* with the PF04326.19 HMM profile from the Pfam database^24^. Prokaryotic Slfn domains (5,930) were extracted and aligned using MAFFT v7.526 with 596 unique mammalian Slfn domain sequences (CD-HIT with `-c 1`) found with *jackhmmer*. The resulting alignment was used to build a phylogenetic tree using Fast-Tree v2.1.11 with `-pseudo -wag -gamma` options. For visualization, the tree was pruned to remove prokaryotic sequences with >50% sequence similarity (CD-HIT -c 0.5) and mammalian sequences with >70% sequence similarity (CD-HIT `-c 0.7`). The resulting tree was visualized and annotated using iTOL^37^.

For every identified *pSlfn*, genes within a *±*10 kbp distance were extracted and annotated using *HMMscan* (i-E-value < 0.001) against HMM profiles downloaded from the DefenseFinder database in March 2025^38^. The number of defense genes identified in the genetic proximity was then calculated for every *pSlfn* gene (Supplementary Table 1). Defense scores were calculated for *pSlfn* clusters as the number of *pSlfn* homologs with at least one annotated defense gene in the genetic neighborhood divided by the total number of *pSlfn* homologs within the cluster.

### Classification of pSlfn proteins

Sequences of 5,930 *pSlfn* proteins were searched against the Pfam database^24^ (v37.2) using HMMER’s *hmmscan*^35^, and against the ECOD database^25^ (v292) using BLAST^39^. High-confidence Pfam hits (i-E-value < 0.0001) were filtered to remove overlapping or nested annotations by selecting the most significant predictions. Proteins with significant sequence homology to multiple Pfam families, i.e., fusions of Slfn (PF04326) to other domains, were further classified by assignment to their respective Pfam clan, representing a higher-order grouping of related domain families. Proteins with Slfn domains and less than 40 amino acids of remaining sequence were annotated as “pSlfn only”. Proteins Slfn domain only and the remaining sequence of more than 40 amino acids (2,134 total) were further annotated using structural homology as described before^40^. Briefly, 2,134 proteins were grouped in 903 clusters using MMseqs with 50% sequence identity and 0.8 coverage cut-offs^41^. AlphaFold 3 (AF3) was used to predict protein structure models for each cluster representative^42^. AF3 models were processed by Domain Parser for AlphaFold Models (DPAM) to identify domain architectures and map domains to the ECOD database classification^43^. The resulting ECOD domain classifications were assigned to cluster members, converted into corresponding Pfam clans (where possible), and integrated with the sequence-based Pfam domain annotation. For 1,020 *pSlfn* proteins, a confident annotation could not be assigned. These proteins were designated as unannotated fusions of *pSlfn* domains and grouped into 558 sequence similarity groups using MMseqs. The domain annotations can be found in Supplementary Table 2.

### Computational search for T5.142 homologs

Homologs of T5.142 were identified in the database of non-redundant protein sequences (NCBI) using PSI-BLAST. Identified homologs were clustered using CD-HIT with -c 0.9, and cluster representatives were used for experimental testing. Protein sequences of T5.142 homologs were aligned with MAFFT v7.526. The phylogenetic tree was constructed using FastTree v2.1.11 with `-pseudo -wag -gamma` options and visualized using iTOL.

### Bacterial growth assays

*E. coli* K-12 MG1655 cells expressing *Ror*Slfn5, *Ror*Slfn5^E15A, D20A^, or carrying no defense system were grown with shaking at 37 °C to an OD_600_ of 0.3. Then, 180 *µ*L of the cell culture was transferred to a 96-well plate and mixed with 20 *µ*L of T5 phage dilution at varying multiplicities of infection (MOI). The 96-wells were shaken and optical density (OD_600_) was measured every 60 seconds for 6 hours at 37 °C using a SpectraMax M5e reader. To measure bacterial growth upon viral trigger expression, *E. coli* K-12 MG1655 cells were co-transformed with *Ror*Slfn5 or *Ror*Slfn5^E15A,D20A^ and a plasmid with an inducible trigger T5.142 or *gfp* gene. Double-transformed cells were plated on LB agar plates with double antibiotic and 0.2% glucose to suppress basal expression. Night cultures of double-transformed cells were used to inoculate liquid cultures, and cells were grown to an OD_600_ of 0.3. After reaching optical density, 180 *µ*L of bacterial cultures were mixed with 20 *µ*L of D-glucose or L-arabinose to a final concentration of 0.2% in a 96-well plate. Optical density and GFP signal were measured every 60 seconds for 6 hours at 37 °C using a SpectraMax M5e reader. All experiments were performed in three or four biological replicates.

### Phage escapers isolation

T5 phage was serially diluted, and 100 *µ*L of each dilution was mixed with 500 *µ*L of *E. coli* K-12 MG1655 cells expressing *Ror*Slfn5 at OD_600_ = 0.3. This mixture was added to 8 mL of soft agar supplemented with ampicillin (100 *µ*g/mL), 10 mM MgCl_2_, and 10 mM CaCl_2_, and overlaid onto LB agar plates containing the same supplements. Plates were incubated overnight at 37 °C. The following day, several individual plaques were picked to inoculate *E. coli* K-12 MG1655 expressing *Ror*Slfn5 (OD_600_ = 0.3). The cultures were incubated overnight in LB media supplemented with ampicillin (100 *µ*g/mL) at 37 °C with shaking at 180 rpm. The next day, cultures were centrifuged at 5,000*×* g for 10 min at 4 °C, and the supernatants containing phages were transferred to fresh tubes. The phages were serially diluted and spotted on the lawns of *E. coli* K-12 MG1655 cells expressing *Ror*Slfn5 or no system to assess their ability to escape the immune system.

### Phage genomic DNA isolation and sequencing

Genomic DNA of T5 phage and T5 phage escapers was isolated from 1 mL of phage supernatant using Norgen Biotek™ Phage DNA Isolation Spin Column Kit (SKU 46800). 500 ng of each genomic DNA sample was used to prepare a sequencing library as described in the SQK-LSK114 protocol using the Native Barcoding Kit 24 V14 (SQK-NBD114.24). The flow cell was primed, and the barcoded sequencing library (300 ng) was loaded according to the Oxford Nanopore protocol (SQK-LSK114). Raw sequencing data (POD5 files) were basecalled in the super-accuracy mode (dna_r10.4.1_e8.2_400bps_sup@v5.0.0) and demultiplexed using *Dorado* basecaller, version 0.8.3 (Oxford Nanopore). Base-called reads were aligned to the reference (NCBI: NC_005859.1) using *minimap2* (v2.28-r1209) with Nanopore preset (-ax map-ont setting). Resulting alignments (BAM files) were used to call consensus sequences using *samtools* v1.21 and call sequence variants using *bcftools* v1.21.

### Cell toxicity assay

*E. coli* K-12 MG1655 cells were co-transformed with plasmids encoding for defense systems and inducible pBAD vectors with viral trigger genes or *gfp*. Double-transformed cells were selected by plating on LB agar supplemented with ampicillin (100 *µ*g/mL), kanamycin (100 *µ*g/mL), and D-glucose (0.2%), followed by overnight incubation at 37 °C. Individual colonies were then used to inoculate overnight cultures in LB medium containing the same antibiotics. The following day, cultures were adjusted to OD_600_ = 0.6, ten-fold serially diluted, and 3 *µ*L of each dilution was spotted onto LB agar plates supplemented with ampicillin (100 *µ*g/mL), kanamycin (100 *µ*g/mL), and either 0.2% D-glucose or 0.2% L-arabinose. Resulting plates were incubated overnight at 37 °C. All experiments were performed in three biological replicates.

### SOS response reporter assay

*E. coli* K-12 MG1655 cells were co-transformed with *Ror*Slfn5 or *Ror*Slfn5^E15A, D20A^ plasmid, pBAD plasmid with T5.142 or *gfp* gene, and pSOS-GFP reporter plasmid. Triple-transformed cells were grown to an OD_600_ of 0.3, and 180 *µ*L of these cultures were mixed with 20 *µ*L of D-glucose or L-arabinose to a final concentration of 0.2% in a 96-well plate. Bacterial growth (OD_600_) and GFP signal were measured every 60 seconds for 6 hours at 37 °C using a SpectraMax M5e reader. Mitomycin C (RPI #M92010) was added to the bacterial culture to a final concentration of 100 nM as a positive control to induce the SOS response. PBS was used as a negative control. All experiments were performed in three biological replicates.

### Total RNA extraction

*E. coli* K-12 MG1655 cells were double-transformed with plasmids expressing either *Ror*Slfn5 or *Ror*Slfn5^E15A, D20A^, along with a plasmid encoding the viral trigger T5.142. Cultures were grown to an OD_600_ of 0.3, and trigger expression was induced by adding L-arabinose to 0.2% final concentration. Cells were collected at 0, 30, and 60 min post-induction for total RNA extraction using the Direct-zol RNA Miniprep Kit (Zymo Research, R2052). Briefly, the cells were spun down at 3,000 *×* g for 5 min at 4 °C. The supernatants were removed, and the cell pellets were resuspended in 900 *µ*L of TRI-reagent (Sigma-Aldrich). Then, the samples in TRI-reagent were loaded on columns, and the total RNA purification was performed according to the kit instructions with a DNAse I on-column treatment step. ~ 500 ng of total RNA sample was used to run a 12% Urea PAGE. The *E. coli* K-12 MG1655 cell cultures expressing *Ror*Slfn5, *Ror*Slfn5^E15A, D20A^, PARIS, or PARIS^AriB(E26A)^ (OD_600_ = 0.3) were infected with T5 phage at an MOI of 10. Culture samples (5 mL) were taken right before the infection (0 min) and 30 min after for total RNA extraction using the Direct-zol RNA Miniprep kit from Zymo Research (R2052) as described above. The quality of RNA samples was assessed using the Agilent 2100 Bioanalyzer at the University of Florida ICBR Gene Expression and Genotyping Core Facility (RRID:SCR_019145).

### RNA sequencing

Total RNA samples (~ 1,200 ng each) were treated with T4 PNK (NEB, M0201) in a reaction buffer (50 mM Tris-HCL, pH 7.5, 10 mM MgCl_2_, 1 mM DTT, 1 mM ATP, and murine RNase inhibitor 1U/ *µ*L (NEB)) for 1 hour at 37 °C. Next, RNA was purified with the Monarch RNA Cleanup Kit (NEB, T2030L). PNK-treated RNA samples were polyadenylated using *E. coli* Poly(A) polymerase (NEB, M0276S) in a reaction buffer (50 mM Tris-HCl, pH 8.0, 250 mM NaCl, 10 mM MgCl_2_, 1 mM ATP, murine RNase inhibitor 1U/ *µ*L) for 30 min at 37 °C. The RNA samples were purified with the Monarch RNA Cleanup Kit (NEB, T2030L) and sonicated with Bioruptor Pico sonication device. For shearing, RNA samples were diluted in TE buffer (10 mM Tris-HCl, pH 8.0, 1 mM EDTA) to 50 *µ*L and sonicated for 30 cycles, with 30 seconds of ON time and 30 seconds of OFF time (15 min total sonication time). RNA was purified with RNAClean XP beads (Beckman Coulter, A63987) using a 2:1 ratio of beads to RNA (v/v) and used to prepare a sequencing library as described in the SQK-PCB114.24 protocol using Barcoding kit CDNA-PCR 24 V14 (Oxford Nanopore). Briefly, the RNA was reverse-transcribed, and the resulting cDNA was amplified with barcoded PCR primers. Barcoded PCR products were purified using Mag-Bind TotalPure NGS magnetic beads (Omega Bio-tek) and pooled together for sequencing library preparation. The flow cell was primed, and the barcoded sequencing library (~ 10 ng) was loaded according to the Oxford Nanopore protocol (SQK-PCB114.24). Raw sequencing data (POD5 files) were basecalled using the super-accuracy model (dna_r10.4.1_e8.2_400bps_sup@v5.0.0) and demultiplexed with *Dorado* basecaller, version 0.9.1, with the `–no-trim` option (Oxford Nanopore). Primer sequences and polyA (sense strand reads) or polyT (anti-sense strand reads) tails were removed using *cutadapt* v5.0. *Bowtie 2* (v2.5.4) was used to align trimmed reads to a reference sequence created by concatenating sequences of the *E. coli* K-12 MG1655 genome (NCBI: NC_000913.3), the T5 phage genome (NCBI: NC_005859.1), and the pAN248 plasmid (this work). To map RNA ends, start and end coordinates for every read were extracted using *bedtools bamtobed* (v2.31.1). RNA end coordinates were mapped to genetic features, quantified, and normalized using total mapped read counts to produce count per million mapped reads (CPM) values. Read alignments were plotted using *Gviz* v1.44.2 R package.

### Protein expression and purification

For individual protein expression and purification, the vectors pAN261 (6xHisTwinStrepSUMO-*Ror*Slfn5) and pAN302 (6xHisTwinStrepSUMO-T5.142) were transformed into *E. coli* BL21(DE3) cells. Cells were grown in LB broth (Lennox) supplemented with ampicillin (100 *µ*g/mL) at 37 °C to an OD_600_ of 0.5-0.7, incubated on ice for 30 min, and then induced with 0.5 mM IPTG for overnight expression at 16 °C. Cells were lysed with sonication in Lysis buffer (20 mM Tris-HCl pH 8.0, 500 mM NaCl, 1 mM TCEP, protease inhibitor (PI78430, ThermoFisher)). Lysates were clarified by sequential centrifugation: first at 10,000 *×* g for 20 min at 4 °C, and then the supernatant was re-centrifuged at 10,000 *×* g for an additional 10 min at 4 °C. The affinity-tagged proteins were bound to StrepTrap XT-columns (Cytiva) and eluted with elution buffer (20 mM Tris-HCl pH 8.0, 500 mM NaCl, 1 mM TCEP, 50 mM Biotin). Proteins were concentrated using 10 kDa (T5.142) or 100 kDa (*Ror*Slfn5) spin concentrators. Affinity tags were removed by overnight dialysis at 4 °C against SUMO digest buffer (30 mM Tris-HCl pH 8.0, 500 mM NaCl, 1 mM TCEP, 0.15% Igepal) with His-tagged SUMO protease made in-house. The affinity tag and protease were removed using a HisTrap HP column (Cytiva), and the flow-through containing untagged *Ror*Slfn5 or T5.142 was concentrated using Corning Spin-X concentrators at 4 °C. Finally, the proteins were purified using a Superdex 200 10/300 size-exclusion column (Cytiva) in a buffer (20 mM Tris-HCl, pH 8.0, 500 mM NaCl, 5 mM MgCl_2_, 1 mM TCEP). Fractions containing the target protein were pooled, concentrated, aliquoted, flash-frozen in liquid nitrogen, and stored at *−*80 °C.

For the purification of the *Ror*Slfn5-T5.142 complex, 6xHisTwinStrepSUMO-tagged *Ror*Slfn5 and untagged T5.142 were expressed separately. *Ror*Slfn5 was expressed as described above. Untagged T5.142 was expressed from plasmid pAN265. *E. coli* BL21(DE3) cells were grown to an OD_600_ of 0.5 and induced with 0.2% L-arabinose overnight at 16 °C. Cell pellets for each protein were lysed with sonication in Lysis buffer (20 mM Tris-HCl pH 8.0, 500 mM NaCl, 1 mM TCEP, protease inhibitor (PI78430, ThermoFisher)) and lysates were clarified by centrifugation as described above. The clarified lysates were combined and incubated at 4 °C for 30 min. The Strep-tagged complex was purified from the combined lysate using a StrepTrap XT column. The affinity tag was removed, and the complex was further purified using a Superdex 200 10/300 size-exclusion column (Cytiva), as described above.

### tRNA synthesis

Bacterial and T5 phage tRNA were synthesized using *in vitro* transcription as previously described^29^. Briefly, single-stranded DNA oligos encoding tRNA with an upstream T7 promoter sequence were ordered from Eurofins Genomics (Supplementary Table 3) and used as templates for PCR. Each 25 *µ*L PCR reaction contained 1X Q5 Reaction Buffer (NEB), 200 *µ*M dNTPs, 0.5 *µ*M forward primer, 0.5 *µ*M reverse primer, 0.2 U/*µ*L Q5 High-Fidelity DNA Polymerase (NEB), and 0.04 *µ*M of DNA oligo template. PCR products were purified using the DNA Clean and Concentrator-5 Kit (Zymo Research, D4014). To synthesize tRNA, 0.8-1 *µ*g of PCR product was used as template for an *In Vitro* Transcription (IVT) reaction using HiScribe® T7 High Yield RNA Synthesis Kit (NEB, E2040L). 10 *µ*L reactions were prepared following the manufacturer’s instructions and incubated for 16-18 hours at 37 °C. IVT products were treated with DNase I (NEB, M0303L) and then purified using the Monarch RNA Cleanup Kit (NEB, T2050S).

### tRNA cleavage assay

To test *Ror*Slfn5 nuclease activity *in vitro*, 100 nM of *Ror*Slfn5 or *Ror*Slfn5^E15A, D20A^ was incubated with 100 nM of tRNA with or without the purified T5.142 trigger (2 *µ*M) in the reaction buffer (50 mM Tris-HCl pH 7.5, 10 mM MnCl_2_ or 10 mM MgCl_2_, 100 mM NaCl_2_, and 1 mM DTT). All reactions were incubated for 1 hour at 37 °C, and 50 mM EDTA was added to stop the reaction. Samples were incubated for 10 min at 70 °C with 2X RNA loading dye before loading into a preheated 15% Urea PAGE. After running the gel at 20W, the gel was stained with SYBR Gold (Invitrogen, S11494) for 30 min and imaged on a Bio-Rad Chemidoc MP Imaging System.

### Statistics & reproducibility

Statistical analyses were performed using R version 4.3.0 with functions from the stats package. Experiments that compare three or more groups were analyzed using one-way Analysis of Variance (ANOVA). Post hoc pairwise mean comparisons were performed using Tukey’s Honestly Significant Difference (HSD) test. Means in experiments with two groups were compared using a two-sided Welch’s t-test. For RNA-seq data, normalized (Counts Per Million – CPM) and log2-transformed RNA end counts were compared using a two-sided Welch’s t-test with unequal variances, and resulting *p* values were adjusted using Benjamini and Hochberg correction to calculate false discovery rate (FDR). The statistical test, levels of statistical significance, and sample size (n) of each experiment are provided in the figure legends.

## Data Availability

Raw sequencing data were deposited in the NCBI Sequence Read Archive under Bioproject accession numbers PRJNA1272589 (phage genome sequencing) and PRJNA1272590 (3^*′*^-end RNA sequencing) and will be released upon publication of the manuscript.

## Code Availability

The code used for RNA-seq data analysis is available upon request.

## Acknowledgements

We thank B. Wiedenheft and R. Wilkinson for sharing *E. coli* strains and bacteriophages; M. Kladde and M. Gauthier for Bioruptor Pico sonicator access; M. Buyukyoruk for advice on computational analyses; University of Florida ICBR Gene Expression and Genotyping Core Facility for RNA analysis using the Agilent 2100 Bioanalyzer (RRID:SCR_019145); A. Nemudryi was supported by a start-up package from the University of Florida College of Medicine, University of Florida Office of the Vice President for Research, and University of Florida Emerging Pathogens Institute. This work was supported by the National Institutes of Health (United States) grant R00AI171893 (A. Nemudryi). Some diagrams and schematics were created with BioRender.com.

## Author contributions

A. Nemudryi and A. Nemudraia conceived, designed, and supervised the project. A. Nemudryi performed the initial bioinformatic search and neighborhood analysis. K.E.M. performed classification of *pSlfn* domain architectures. A. Nemudryi, A. Nemudraia, R.C., and V.P.T. designed and generated the plasmids. A. Nemudraia and R.C. performed phage plaque assays. A. Nemudryi performed a bacterial growth assay. A. Nemudraia and A. Nemudryi performed phage escapers isolation and sequencing. A. Nemudryi and V.P.T. performed the cell toxicity assays. A. Nemudryi and A. Nemudraia performed RNA extractions and sequencing experiments. A. Nemudryi, A. Nemudraia, Y.W., and L.L. performed protein expression and purification. V.P.T. performed tRNA synthesis and tRNA cleavage assay. A. Nemudryi and A. Nemudraia wrote the original draft, with input from all authors.

