## Supplementary Figures for "Bacterial Schlafens mediate anti-phage defense"

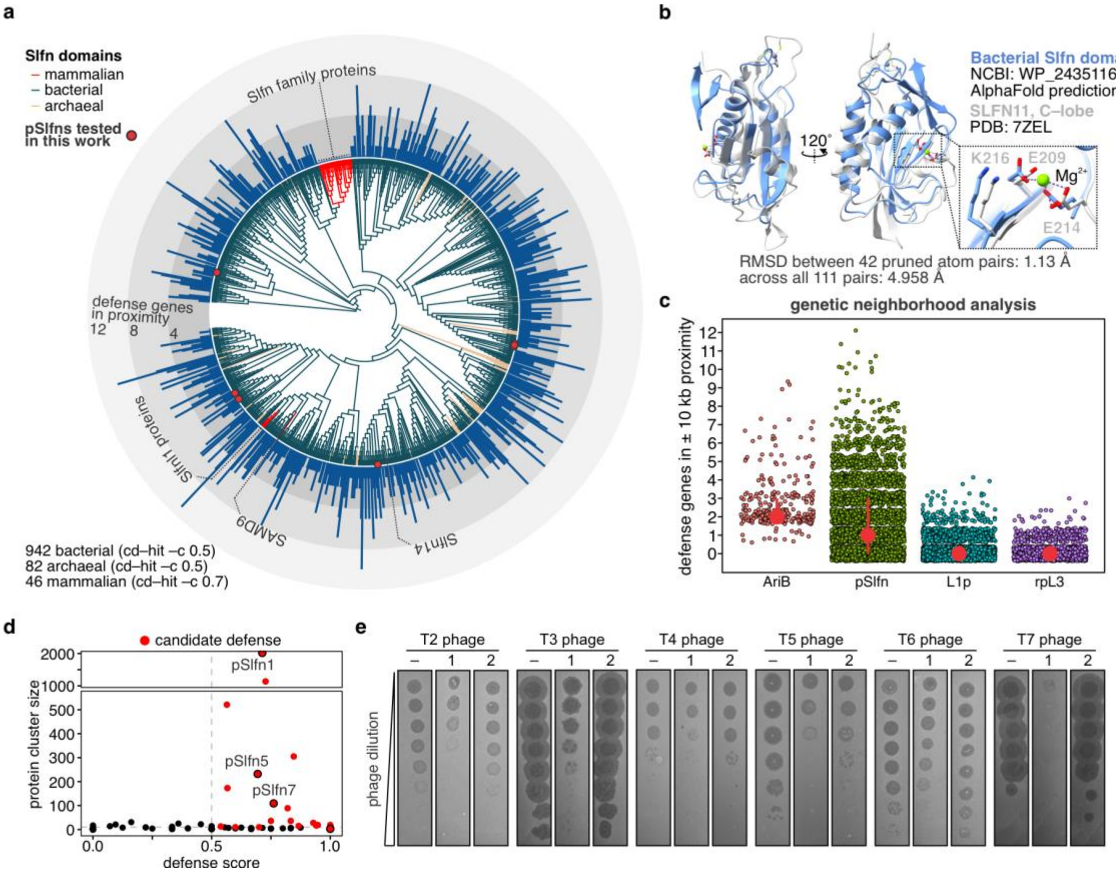


**Supplementary Fig. 1. Computational analysis of prokaryotic Schlafen (Slfn) domains. a,** phylogenetic tree of bacterial, archaeal, and mammalian Slfn domains. Tree branches are colored according to the taxonomic group. To improve visualization, the phylogenetic tree was pruned to keep representative sequences identified by CD-HIT from the original dataset. A 50% CD-HIT threshold was used for prokaryotic Slfn domains, while mammalian Slfn domains were pruned with a 70% threshold. Blue bars show the number of phage defense genes in the ±10 kb neighborhood of prokaryotic Slfn genes. **b,** alignment of the AlphaFold predicted structure of the bacterial Slfn domain (light blue) and the C-lobe of the human SLFN11 nuclease domain. The inset shows the ribonuclease active site with amino acid residues required for the catalytic activity. **c,** Scatter plot showing results of genetic neighborhood analysis for AriB (known anti-phage gene), pSlfn, L1p, and rpL3 genes. L1p and rpL3 were used as controls with established non-immune functions. Each dot represents the number of known phage defense genes within a 10 kb neighborhood of AriB, pSlfn, L1p, or rpL3 homologs in our database of prokaryotic genomes. The red dot shows the median value. Vertical red lines show interquartile ranges. **d,** pSlfn proteins were clustered based on domain architecture annotated using sequence and structural homology methods. Each point represents a pSlfn protein group. *Y-axis:* number of unique sequences in the group; *x-axis*: defense score, which was calculated as the ratio of group members that are found within ± 10 kbp distance from at least one known defense gene to the total number of mebers in the group (see Methods). Clusters with more than ten representatives (horizontal dashed line) and a defense score > 0.5 (vertical dashed line) were further considered as candidate defense systems (red dots). Black circles show domain architectures experimentally tested in Figure 1. **e,** A representative phage spot assay showing that *Eco*DUF262-Slfn (1) and *Ror*Slfn5 (2) protect from phages.

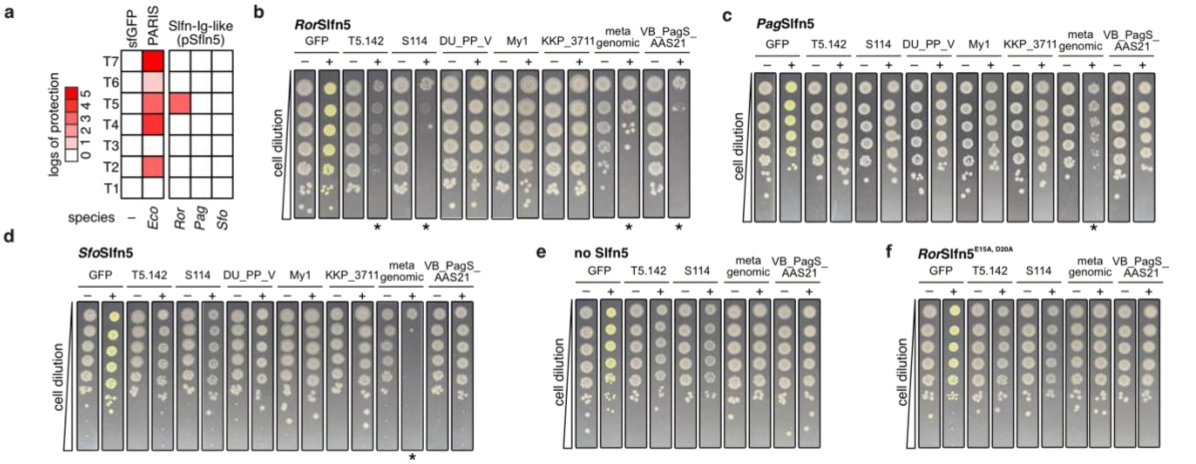


**Supplementary Fig. 2.** ***T5 phage tail assembly protein triggers pSlfn5-mediated abortive infection phenotype.*** **a,** Antiviral activities of pSlfn5 homologs (left-to-right) against a panel of phages (top-to-bottom). **b-d,** Cellular toxicity assay in *E. coli* K-12 MG1655 cells co-transformed with a plasmid expressing *Ror*Slfn5 (**b**), *Pag*Slfn5 (**c**), or *Sfo*Slfn5 (**d**) and a plasmid for arabinose-inducible expression of T5.142 homologs (T5.142 – T5 phage, S114 – Salmonella phage S114, DU_PP_V - Pectobacterium phage DU_PP_V, My1 - Pectobacterium phage My1, KKP_3711 - Enterobacter phage KKP_3711, vB_PagS_AAS21 - Pantoea phage vB_PagS_AAS21). Asterisk (*) indicates toxicity phenotypes. pBAD-GFP was used as a positive control for inducible protein expression. **e-f**, Cellular toxicity assay in MG1655 transformed only with T5.142 homologs (**e**) or co-transformed with a plasmid expressing *Ror*Slfn5$^{E15A,D20A}$ and T5.142 homologs (**f**).

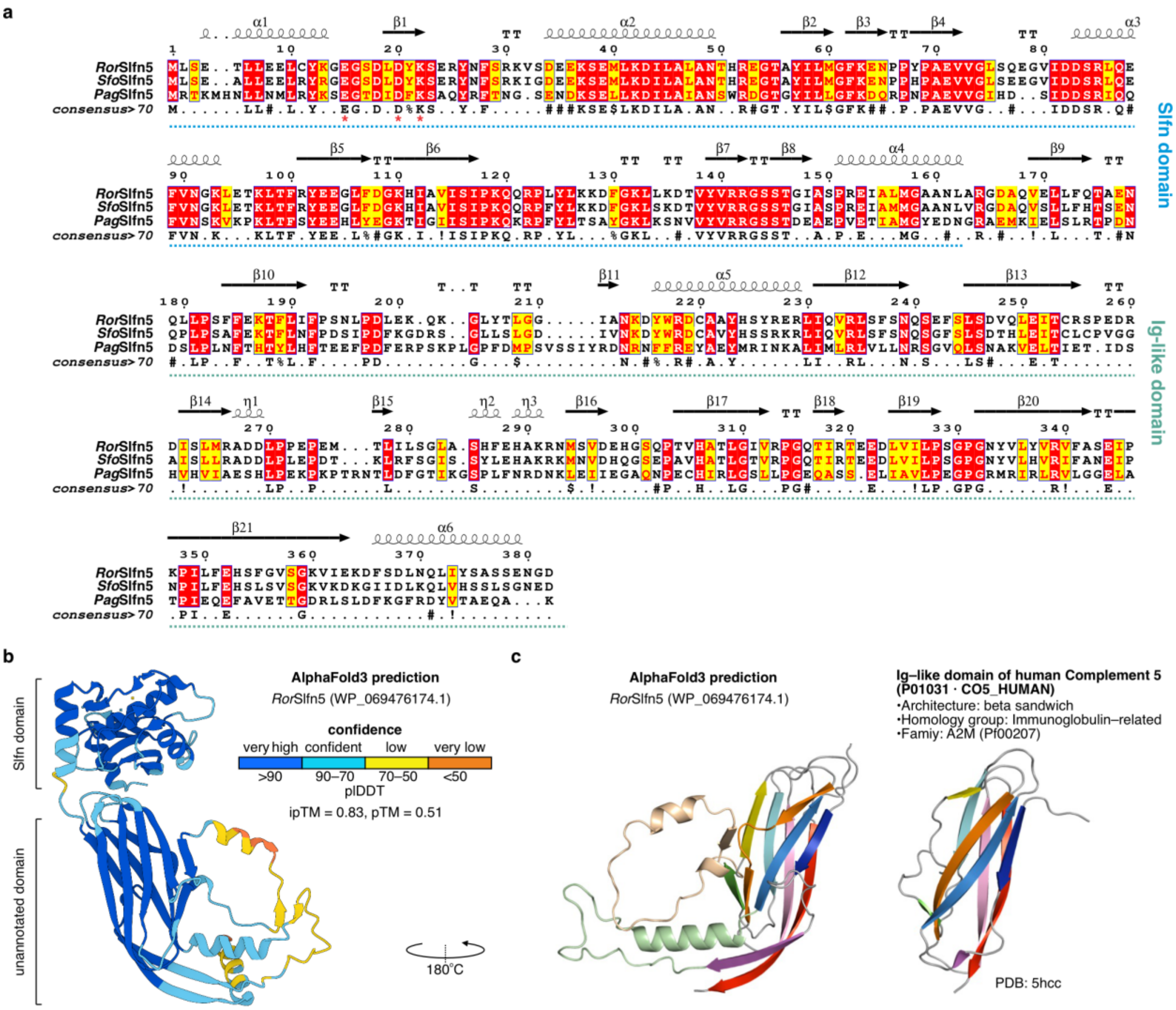


**Supplementary Fig. 3.** The **C-terminus of pSlfn5 contains an Ig-like phage sensor domain. a,** Multiple sequence alignment of pSlfn5 homologs. Red asterisks (*) indicate the conserved catalytic motif of the Slfn ribonuclease domain. **b,** AlphaFold-predicted structure of *Ror*Slfn5. **c,** Comparison of AlphaFold-predicted structures of the C-terminal domain of RorSlfn5 and the Ig-like domain of human Complement 5.

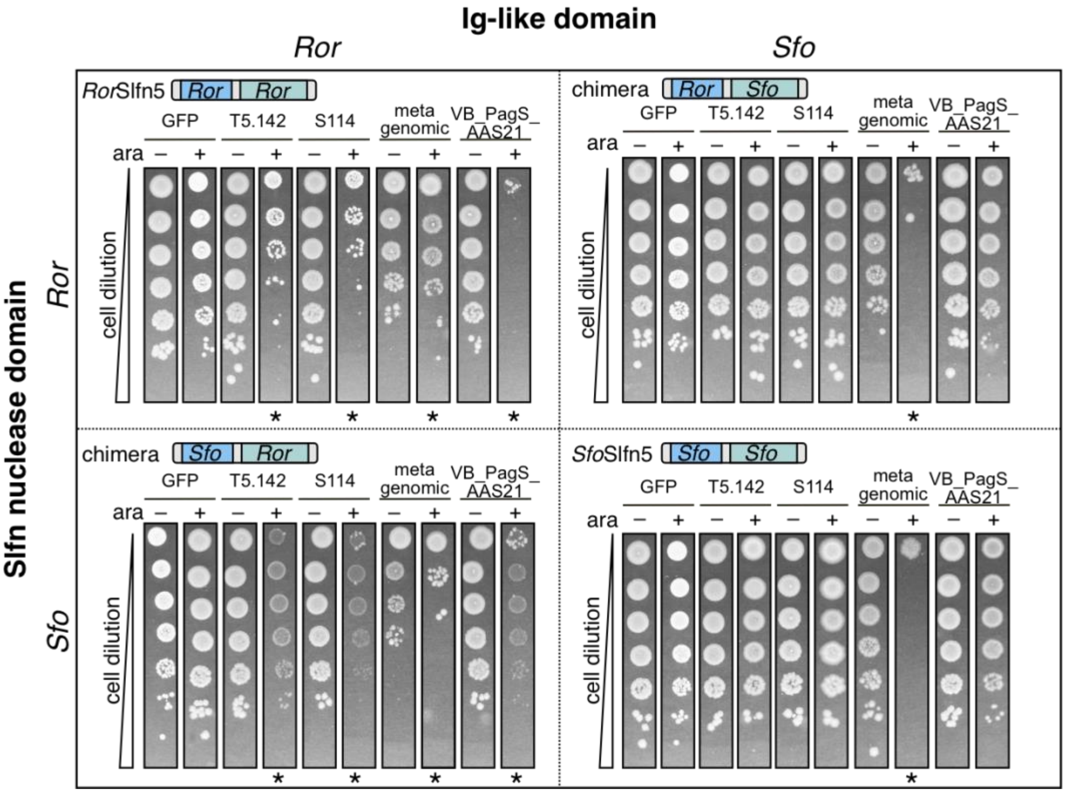


**Supplementary Fig. 4. Ig-like domain of pSlfn5 defense confers phage trigger specificity.** Cellular toxicity assay in E. coli K-12 MG1655 cells co-transformed with a plasmid expressing *Ror*Slfn5, *Sfo*Slfn5*,* or chimeric proteins with swapped Ig-like domains and a plasmid for arabinose-inducible expression of T5.142 homologs (T5.142 – T5 phage, S114 – Salmonella phage S114, DU_PP_V - Pectobacterium phage DU_PP_V, My1 - Pectobacterium phage My1, KKP_3711 - Enterobacter phage KKP_3711, vB_PagS_AAS21 - Pantoea phage vB_PagS_AAS21). Asterisk (*) indicates toxicity phenotypes. pBAD-GFP was used as a positive control for inducible protein expression.

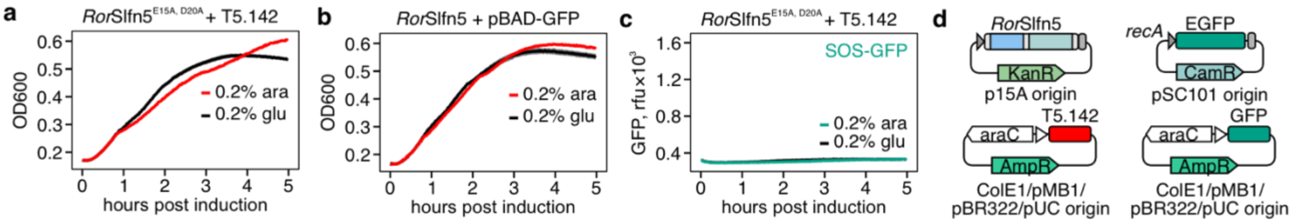


**Supplementary Fig. 5. Additional controls for main Fig. 4. a,** growth kinetics of MG1655 cells co-transformed with *Ror*Slfn5 and inducible T5.142 plasmid after addition of L-arabinose or D-glucose. **b,** Same for MG1655 co-transformed with *Ror*Slfn5 plasmid and pBAD-GFP. **c,** SOS response reporter assay for MG1655 cells co-transformed with plasmid encoding inactivated *Ror*Slfn5 defense (E15A, D20A mutation). **d,** Schematics of different plasmids used in the SOS response and cell toxicity assays.

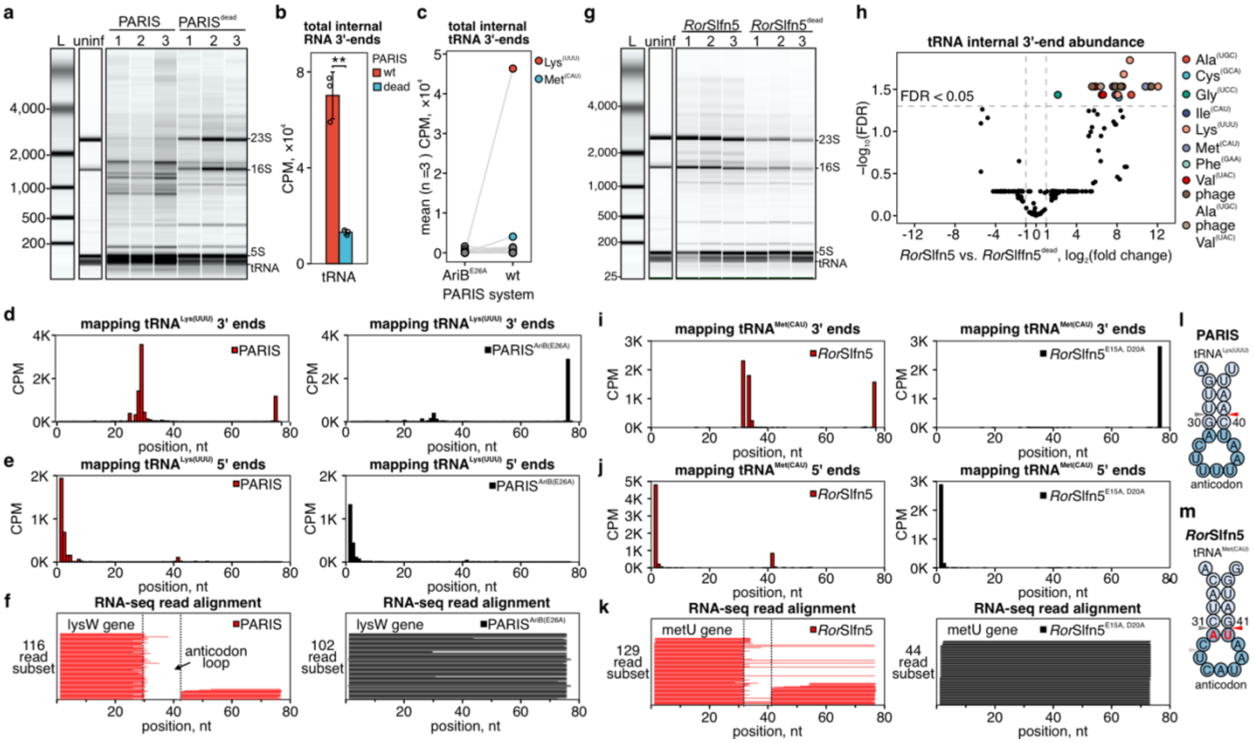


**Supplementary Fig. 6. Phage-activated *Ror*Slfn5 cleaves tRNA anticodon arm. a,** Agilent 2100 Bioanalyzer gel image for total RNA extracted 30 min post T5 phage infection of MG1655 expressing active PARIS defense or its inactive mutant [PARIS$^{dead}$; AriB(E26A) mutation]. 1,2,3 – biological replicates. Red asterisks mark phage-associated transcripts absent in the uninfected (uninf) control. **b,** quantification of total 3'end abundance in tRNA from phage-infected cells expressing PARIS defense vs. PARIS$^{dead}$ control. Data is shown as the mean of three biological replicates ± S.D. Means were compared using a two-sided Welch's t-test. ***p* < 0.01 .**c,** quantification of total 3'-ends counts in specific tRNAs. Data is shown as the mean of three biological replicates. **d-e,** position-specific mapping of 3'-ends (**d**) and 5'-ends (**e**) in tRNA$^{Lys(UUU)}$ in cells with active (left) or inactive (right) PARIS defense. Data is shown as the mean of three biological replicates. **f,** alignment of sequencing reads to the lysW gene of MG1655 cells in cells with active (left) or inactive (right) PARIS defense. One representative replicate of three biological replicates is shown. **g,** Agilent 2100 Bioanalyzer gel image for total RNA extracted 30 min post T5 phage infection of MG1655 expressing active *Ror*Slfn5 defense or its inactive mutant (*Ror*Slfn$^{E15A,\ D20A}$). 1,2,3 – biological replicates. Red asterisks mark phage-associated transcripts absent in the uninfected (uninf) control. **h,** position-specific quantification of internal tRNA 3'-ends in cells expressing active vs. inactive *Ror*Slfn5 defense. FDR – false discovery rate. **i-j,** position-specific mapping of 3'-ends (**i**) and 5'-ends (**j**) in tRNA$^{Met(CAU)}$ in cells with active (left) or inactive (right) *Ror*Slfn5 defense. Data is shown as the mean of three biological replicates. **f,** alignment of sequencing reads to the metU gene of MG1655 cells in cells with active (left) or inactive (right) *Ror*Slfn5 defense. One representative replicate of three biological replicates is shown. **l,** position of PARIS nuclease-dependent RNA ends in tRNA$^{Lys(UUU)}$. The red triangle shows the position of mapped internal 5'-ends; the Gray triangle shows the position of mapped internal 3'-ends. **m,** same for *Ror*Slfn5 nuclease-dependent RNA ends in tRNA$^{Met(CAU)}$.

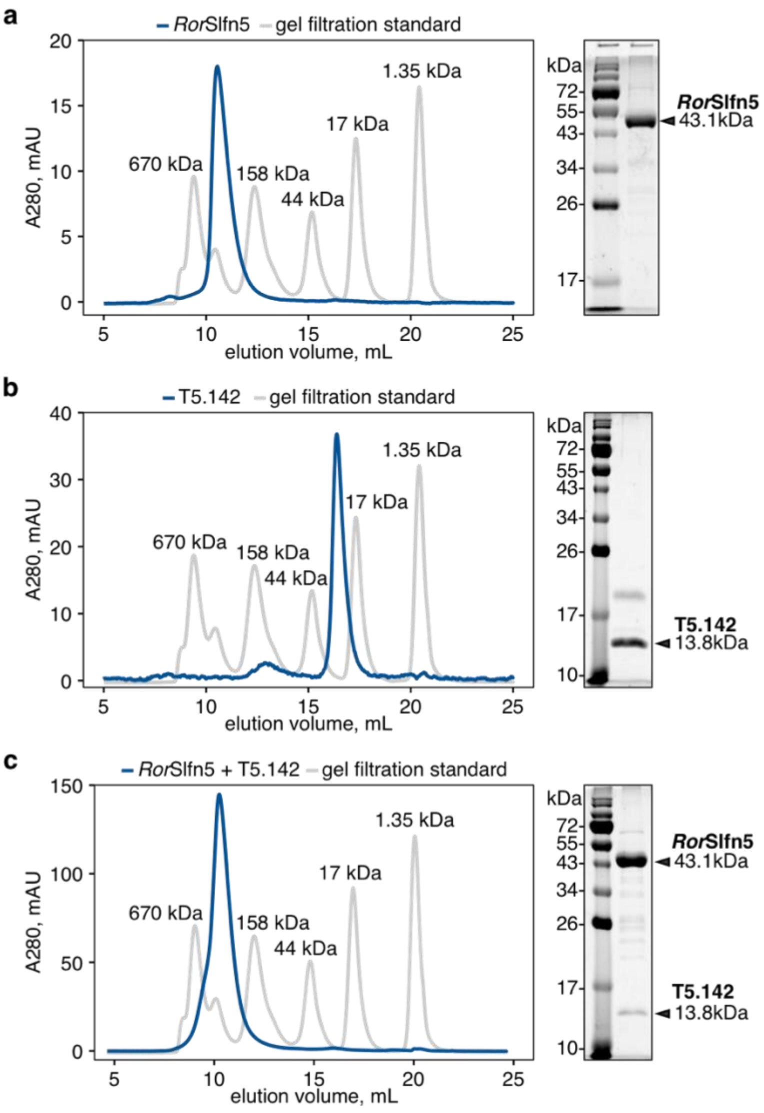


**Supplementary Fig. 7. *Ror*Slfn5 and T5.142 form a stable complex.** Size-exclusion chromatography (SEC) profiles of *Ror*Slfn5 (**a**) and T5.142 (**b**) proteins purified separately or co-purified together (**c**) on a Superdex 200 10/300 GL column. Gel images on the right show SDS-PAGE of SEC peak fractions.

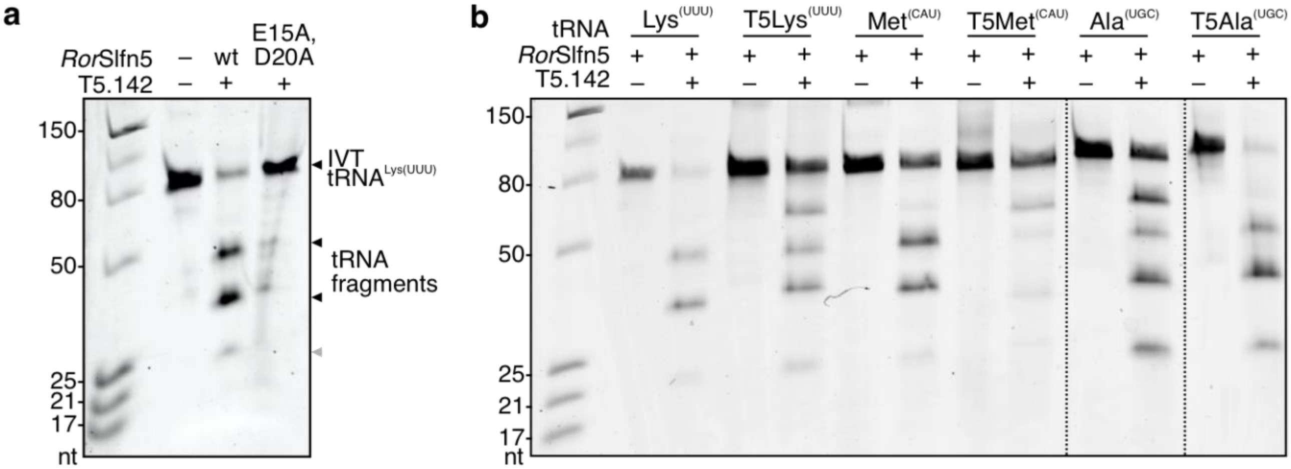


**Supplementary Fig. 8. T5.142 triggers *Ror*Slfn5-mediated tRNA cleavage. a,** tRNA cleavage assays with 100 nM $tRNA^{Lys(UUU)}$ and 100 nM RorSlfn5 or $RorSlfn5^{E15A,\ D20A}$ in the presence of trigger T5.142 (2 µM). RNA fragments were resolved with 15% Urea-PAGE. **b,** Urea-PAGE of 100 nM IVT bacterial and phage (T5) tRNAs after incubating with 100 nM of *Ror*Slfn5 with or without the trigger T5.142 (2 µM).
